# CV-α: designing validations sets to increase the precision and enable multiple comparison tests in genomic prediction

**DOI:** 10.1101/2020.11.11.376343

**Authors:** Rafael Massahiro Yassue, José Felipe Gonzaga Sabadin, Giovanni Galli, Filipe Couto Alves, Roberto Fritsche-Neto

## Abstract

Usually, the comparison among genomic prediction models is based on validation schemes as Repeated Random Subsampling (RRS) or K-fold cross-validation. Nevertheless, the design of training and validation sets has a high effect on the way and subjectiveness that we compare models. Those procedures cited above have an overlap across replicates that might cause an overestimated estimate and lack of residuals independence due to resampling issues and might cause less accurate results. Furthermore, posthoc tests, such as ANOVA, are not recommended due to assumption unfulfilled regarding residuals independence. Thus, we propose a new way to sample observations to build training and validation sets based on cross-validation alpha-based design (CV-α). The CV-α was meant to create several scenarios of validation (replicates x folds), regardless of the number of treatments. Using CV-α, the number of genotypes in the same fold across replicates was much lower than K-fold, indicating higher residual independence. Therefore, based on the CV-α results, as proof of concept, via ANOVA, we could compare the proposed methodology to RRS and K-fold, applying four genomic prediction models with a simulated and real dataset. Concerning the predictive ability and bias, all validation methods showed similar performance. However, regarding the mean squared error and coefficient of variation, the CV-α method presented the best performance under the evaluated scenarios. Moreover, as it has no additional cost nor complexity, it is more reliable and allows the use of non-subjective methods to compare models and factors. Therefore, CV-α can be considered a more precise validation methodology for model selection.

## Introduction

Genomic prediction (GP) proposed by Meuwissen et al. (2001) evolved over the years, but it aims to estimate breeding values of unevaluated genotypes. Hence, it is an important tool for plant breeders to shorten the breeding cycle, increase selection accuracy, and assess genetic variation (Heff et al. 2010; Crossa et al. 2017). Usually, to evaluate the prediction accuracy of the genomic prediction models, the data is divided into training and validation sets. The first set is used to fit the genomic prediction model and estimate the marker effects, whereas the validation set is used to validate the effects estimated in the training set and estimate the accuracy of the predictions (Crossa et al. 2011).

In the genomic prediction context, several methods and parameters have been proposed for the comparison of prediction models (Blondel et al. 2015). Nevertheless, the predictive ability and the bias of the measures are two of the most commonly utilized to evaluate the superiority and goodness of models and scenarios. The former is estimated by Pearson’s correlation between the predicted and true breeding values of individuals contained in the validation set. The latter is obtained by regressing the predicted breeding values over the true ones to obtain the regression coefficient, which indicates the shrinkage (compression) between both (Piepho et al. 2008; Luan et al. 2009).

Some studies have shown that the model accuracy is influenced by the training and validation set (Akdemir et al. 2015; Wu et al. 2015; Auinger et al. 2016), being the main schemes to design training and validation sets in GP studies are K-fold cross-validation (Burgueño et al. 2012; Crossa et al. 2014; Fè et al. 2016) and Repeated Random Subsampling (RRS), also called Monte Carlo CV (Würschum et al. 2014; Yu et al. 2016; Zhang et al. 2016). The first consists of splitting the data into k groups (folds) and fit a model using each fold as training and validation sets. Is this sense, if k = 5, the model will be fitted five times. The second consists of randomly split the dataset into training and validation sets. Both schemes are generally repeated *n* times (see Arlot and Celisse, 2010).

The accuracy estimate obtained by K-fold might be affected by the number of folds, fold size, and the number of replicates (Wong 2015). Likewise, in cross-validation schemes, the RRS is influenced by the relation between training and validation sets and the number of replicates (Kohavi 1995). Furthermore, some factors may lead to biased estimates of predictive ability, such as overlapping between the training and validation set and different relatedness between individuals through sets (Runcie e Cheng 2019). The overlap between training and validation sets over replicates may cause biased results due to the predictions be correlated and non-independent residuals (Amer e Banos 2010). Therefore, neither validation schemes guarantee independence among replicates due to resampling issues. Thus, researchers cannot use standard and non-subjective methods to compare models and factors, such as ANOVA and other multiple comparison tests, due to assumptions unfulfilled regarding residuals independence.

It is import point out that as the number of treatments increases, it becomes a challenge to design orthogonal training and validation sets across the replicates without increase substantially the number of replicates. This problem is similar to experimental field designs involving a large number of treatments. However, the balanced incomplete blocks design seeks to maintain homogeneity among blocks and orthogonality across replicates (Yates, 1936). These schemes are widely used to evaluate the quality of models and their selection for field experiments. Moreover, an extension of cross-validation (CV) schemes applying balanced incomplete block design was first proposed by Shao (1993), considering that each fold is treated as “block” and each genotype as a “treatment.” The orthogonal distribution of the treatments across the blocks within replicates in the balanced incomplete block designs will guarantee that every pair of treatments appears together according to some rules. Therefore, the CV schemes using the incomplete block design may increase the quality of estimates (Fuchs e Krautenbacher 2016), residuals independence, and may allow further multiple comparison analyses.

Based on described above, in this study, we propose a new method to design the training and validation sets for genomic prediction studies based on an alpha-lattice design scheme, called cross-validation alpha-based design (CV-α) and compare its performance to the methods commonly applied in GP studies for model selection. Also, based on the CV-α results, a case of study, via analysis of variance (ANOVA), we could compare the proposed methodology to RRS and K-fold, applying four genomic prediction models with a simulated and real dataset.

## Material and Methods

In order to demonstrate the properties of the proposed cross-validation scheme, we aimed to mimic a standard genomic prediction study, for instance, comparing kernels and statistical methods. Thus, our aim is not comparing genomic matrices or Bayesian and frequentists approaches but simply show that our cross-validation scheme allows multiple comparison tests. For that, we create a simulated population (knowing the true parameters) and also used a well-known real dataset.

### Simulated dataset

We simulated a population of maize single-crosses from inbred parents to perform genomic prediction studies. For this, we used the *AlphaSimR* package (Gaynor 2019). A founder population of 1,000 individuals was simulated with ten chromosomes containing 30,000 segregating loci (SNPs). The individuals were inbred and diploid. Forty-nine individuals were randomly sampled and crossed to compose a partial diallel to obtain 906 hybrids. The phenotypic value (adjusted mean based on heritability) was simulated by randomly sampling 500 QTN from the segregating loci with mean 100 and variance 50. The narrow and broad-sense heritabilities were set to be equal to 0.30 and 0.50, respectively. Finally, to understand the effect of the validation methods in the predictive ability and bias of the true genetic (TGV) and phenotypic value, we performed genomics prediction using both metrics. We repeated the simulations 25 times and averaged the estimates above.

### An empirical case of study: USP maize dataset

We used a dataset of 906 maize single-crosses from a full diallel among 49 tropical inbred lines, according to Griffing’s method 4 (Griffing 1956). The experiments were evaluated in two locations, two years, and under two nitrogen levels. The genotypic information from the 49 tropical inbred lines was obtained from Affymetrix^®^ Axiom^®^ Maize Genotyping Array, containing about 614,000 SNPs (Unterseer et al. 2014). For more details about the phenotypic and genotypic data, see (Fristche-Neto et al. 2018).

The markers with a lower call rate (< 95%), heterozygous loci on at least one individual, and linkage disequilibrium (> 0.90) were removed. The missing markers were imputed using the *Beagle 4*.*0* algorithm (Browning e Browning 2009) from the *synbreed* R package (Wimmer et al. 2012). Later, the genotype of each hybrid was built by combining the genotypes of its parental lines and hybrids with minor allele frequency (MAF < 0.05) were removed. After quality control, a total of 32,207 SNPs was available for further analysis.

To perform the genomic prediction studies, we evaluated the grain yield (GY, Mg ha^-1^), corrected to 13% moisture, and stand across the eight environments. It was used to estimate the BLUP for hybrids following the model:

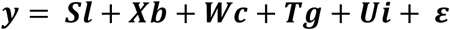

where **y** is the vector of the phenotypic value of hybrids; **l** is the vector of fixed effects of the environment (the combination of site x year x N level); **b** is the vector of fixed effects of blocks within an environment; **c** is the vector of fixed effects of checks; **g** is genotypic values, where 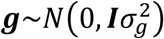 **i** is the interaction between environments and checks, where 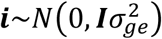; **ε** is the vector of random residuals from checks and hybrid by environments effects, where 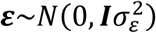. 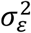 was jointly estimated based on *e* environments with *r* replicated checks in each site. **S, X, W, T**, and **U** are the incidence matrices for **l, b, c, g**, and **i** (Fristche-Neto et al. 2018).

### Genomic prediction

To perform the genomic prediction, we used the additive GBLUP model and the Reproducing Kernel Hilbert Spaces regression (RKHS). The following model equation is the general form of these two approaches:

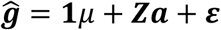

where ***ĝ*** is the vector of BLUP; μ is the intercept; **a** is the vector of additive genetic effects with 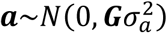; and **ε** is the vector of random residuals with 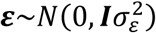. **1** is the incidence vector of μ, and **Z** is the incidence matrix for ***a***. **G** is the genomic relationship matrix (**G**_**a**_ – additive genomic relationship matrix, and **K** – for Gaussian kernel), and **I** is the identity matrix. 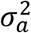 is the additive genetic variance for **G**_**a**_ or genetic variance for **K**, and 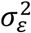 is the residual variance. The additive genomic relationship matrix (**G**_**a**_) was calculated as 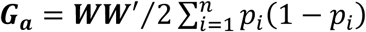, where **W** is the centered matrix of SNPs, and *p*_*i*_ is the frequency of the allele *p* in locus *i* (VanRaden 2008). The Gaussian kernel (**K**) was calculated as 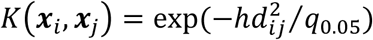, where **x**_**i**_ and **x**_**j**_ are the marker vectors for the i^th^ and j^th^ individuals, respectively, and q_0.05_ is the fifth percentile for the squared Euclidean distance 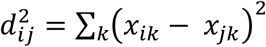 (Pérez-Elizalbe et al., 2015). The *h* value was considered equal to 1.

We used fitted two prediction models considering two statistical approaches (frequentist and Bayesian Genomic Best Linear Unbiased Predictor - GBLUP), resulting in four scenarios: 1) GBLUP with **G**_**a**_ kernel (GA_MM); 2) GBLUP with **K** kernel (GK_MM); 3) Bayesian GBLUP with **G**_**a**_ kernel (GA_Bayes), and 4) Bayesian GBLUP with **K** kernel (GK_Bayes).

The analyses were performed using *ASReml-R* (Butler et al. 2009), and *BGLR* (Pérez and de los Campos, 2014) packages for R. For Bayesian GBLUP models were performed using 10,000 iterations, 3,000 burn-in, and 5 thinning values. The convergence checks for Bayesian models are available in the Supplemental Figure S1 and S2.

### Cross-validation alpha-based design

The cross-validation alpha-based design (CV-α) is an extension of the methodology presented by (Shao 1993) and consists of assigning treatments to folds in each replication by applying the alpha-lattice sorting premises. The CV-α was intended to create scenarios with two, three, or four replicates, regardless of the number of treatments. Each replicate is split into folds, and the number of folds will determine the percentage of training and validation sets. Each fold across replicates is based on the α(0,1) lattice design aiming to reduce the concurrences of any two treatments in the same fold (block) across the replicates (Patterson e Williams 1976).

However, the α(0,1)-lattice design assumptions involve the number of blocks (*s*) and block size (*k*) (number of the folds and fold size, in our context) to determine the number of treatments (Patterson e Williams 1976). As the number of treatments is variable in a real scenario, we compute the nearest smallest number to attend the assumptions above, and the remaining treatments are randomly allocated into the folds. The alpha lattice design was created using the *agricolae* package (Mendiburu 2019), and the scripts are available at Github (https://github.com/allogamous/CV-Alpha).

In order to compare CV-α with the other two benchmarks schemes, we simulated two scenarios: 5-folds with four replicates and 10-folds with two replicates. First, we simulated a scenario with the number of treatments varying from 200 to 2,000 and computed the percentage of remaining treatments that were randomly assigned into folds for each scenario. After, we compared the same two benchmarks schemes according to the mean and standard deviation of the concurrence of any two genotypes, i.e., the number of folds containing both genotypes. The simulations were replicated ten times.

### Model comparison

To evaluate the cross-validation alpha-based design (CV-α) performance, we compared it to benchmark validation schemes: repeated random subsampling (RRS) and K-fold using real and simulated datasets for genomic prediction. For RRS, we used 100 replicates, each with 80% of the data for the training set, and the remaining 20% of the data for the validation set, whereas for CV-α and K-fold were used five-folds and four replicates. The number of replicates or folds for each method considers the most common values for genome prediction studies using Bayesian and frequentist approaches (Zhao et al. 2013; Zhang et al. 2015, 2016; Yu et al. 2016).

From those, we obtained the predictive ability of each statistical model for the different CV methods. The predictive ability was estimated as Pearson’s correlations between the predicted and observed phenotypes. For each CV method, we estimated the slope coefficient for the regression of the predicted values of the validation sets on its phenotypes. For this, the regression coefficient between predicted and genetic value was considered the prediction bias, measuring the degree of inflation/deflation of prediction genomics. Nonbiased models are expected to have a regression coefficient equal to 1. For CV-α and K-fold, the level of averaging considered was at replicates. Although RRS and K-fold schemes do not have independence between replicates, ANOVA have been used to compare the predictive abilities from different models, even breaking the independence assumption. To verify how variance components of models are affected by these methods, we perform the ANOVA test considering the following model:

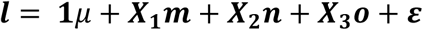

where ***l*** is the vector of Pearson correlation transformed by Fisher z-transformation using the R package *DescTools* (Signorell et al., 2019); μ is the overall mean; ***m*** is the vector of statistical approach effect; ***n*** is the vector of relationship kernel; **o** is the vector of interaction between statistical approach and kernel; and **ε** is the vector of residuals. **X**_**1**_, **X**_**2**_ e **X**_**3**_ are incidence matrices for ***m, n***, and ***o***, respectively. Quadratic components were estimated by the method of moments based on mean square expectation.

## Results

### CV-α

We performed several analyses to evaluate cross-validation alpha-based design (CV-α) performance. For this, we computed the number of treatments that were randomly assigned among folds and the concurrence between pairs of treatments in the same fold across replicates (Figure 1). The results reveal that the proportion of treatments randomly assigned among folds reduces as the number of treatments increases and tends to converge to 0.38% and 0.27% for five-folds with four replicates and ten folds with two replicates, respectively (Figure 1). Considering the concurrence between pairs of treatments in the same fold across replicates, the CV-α reveals lower mean and standard deviation in both evaluated scenarios when compared with the K-fold CV (Figure 2).

**Figure 1.**
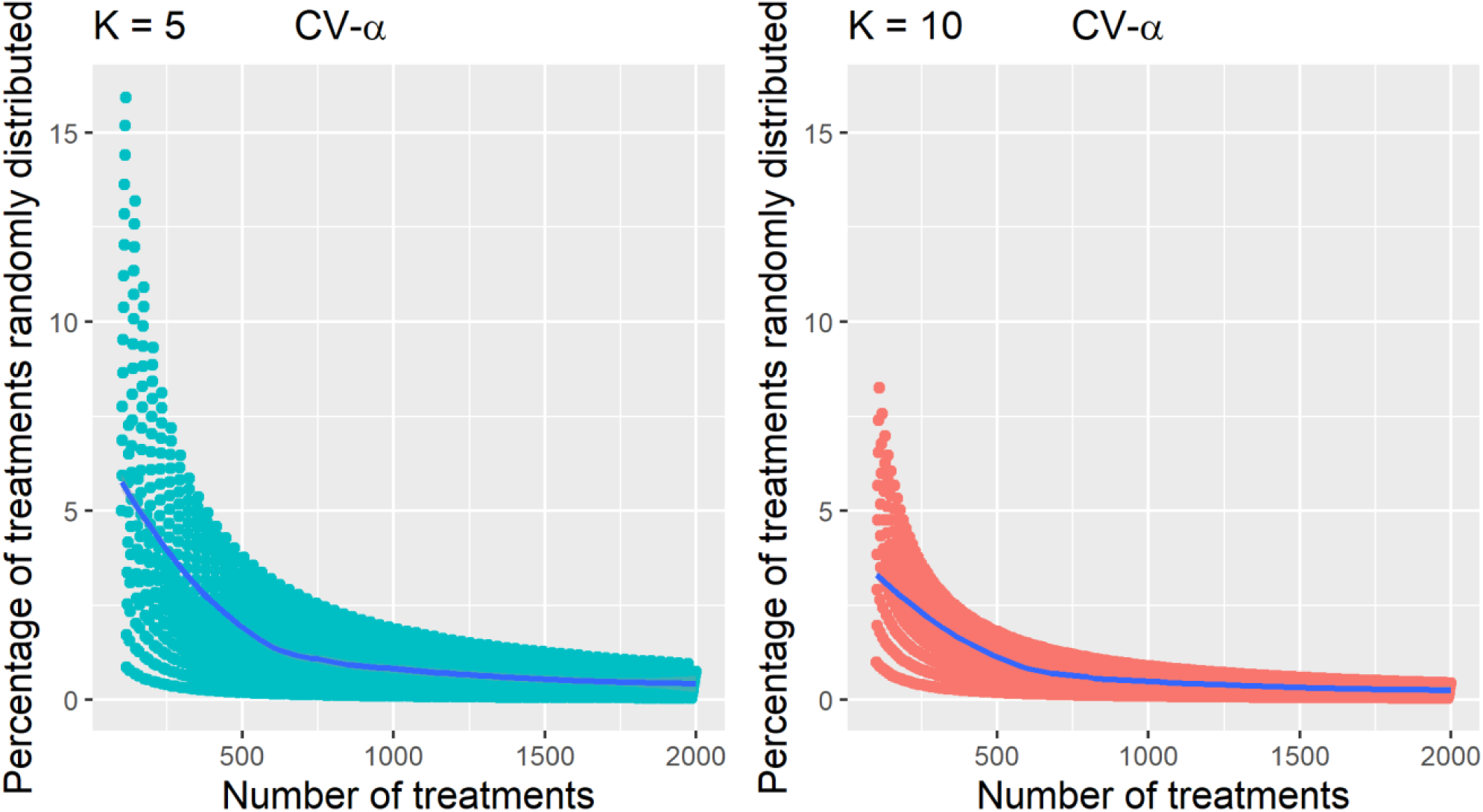
The proportion of treatments randomly distributed into folds to attend the alpha-design presupposition using CV-α with 5-folds with four replicates (a), and 10-folds with two replicates (b), based on simulated data.

**Figure 2.**
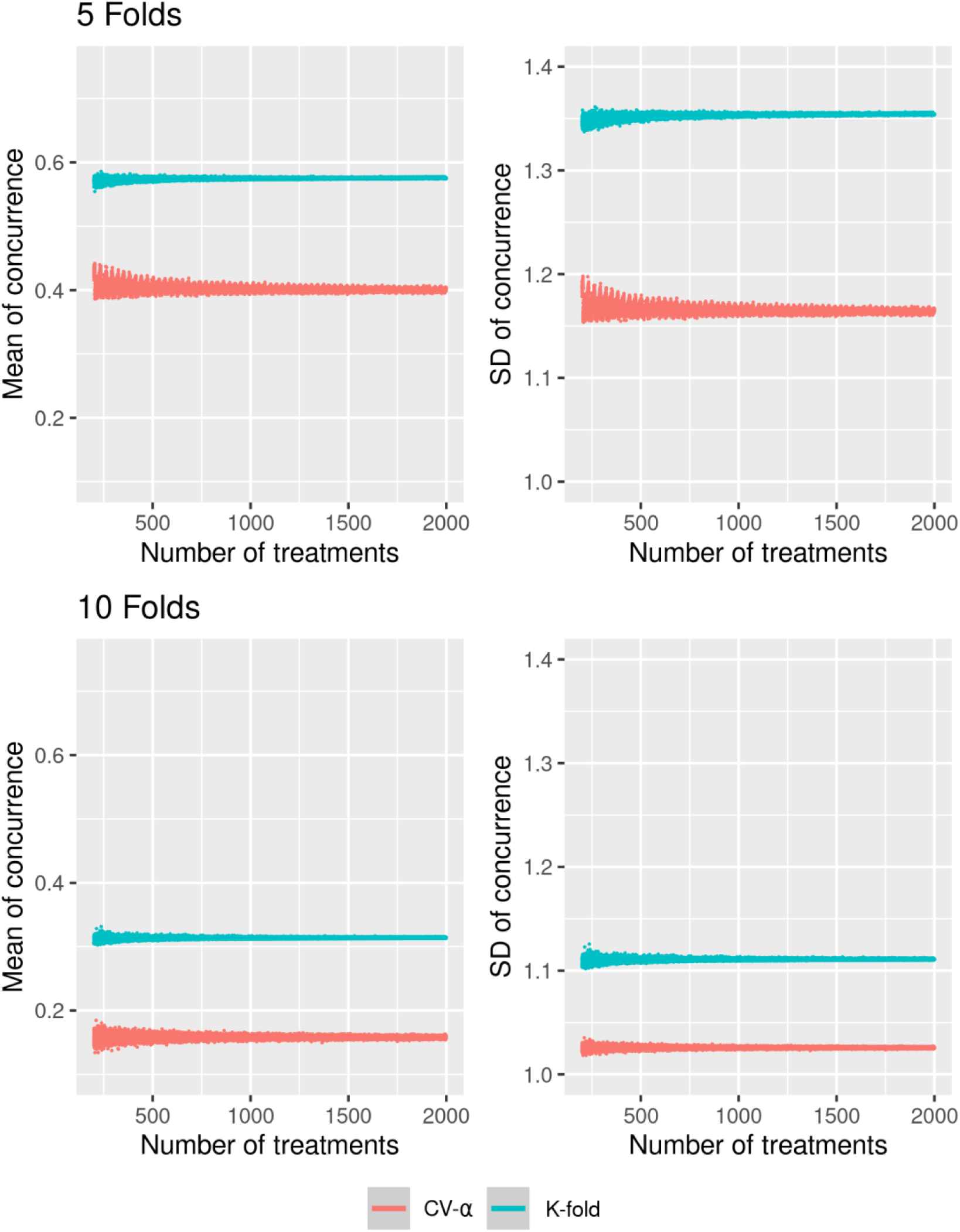
Concurrence (number of times that a pair of treatments appear together in the same fold) mean and standard deviation between treatments pairs in the same fold across replicates using CV-α and K-fold with 5 and 10 folds with 4 and 2 replicates, respectively, based on simulated data.

### Genomic prediction (simulated dataset)

To understand the effects of validation schemes on genomic prediction, we simulated populations to obtain true genetic values (TGV) and phenotypic values. The validation methods did not significantly influence the average prediction ability of TGV and phenotypic values. Nevertheless, the RRS has several “extreme” values when compared to K-fold and CV-α. Besides, RRS showed a more substantial variation for bias, with several values overtaking 0.5 and 1.5 for phenotypic and TGV. (Figure 3).

**Figure 3.**
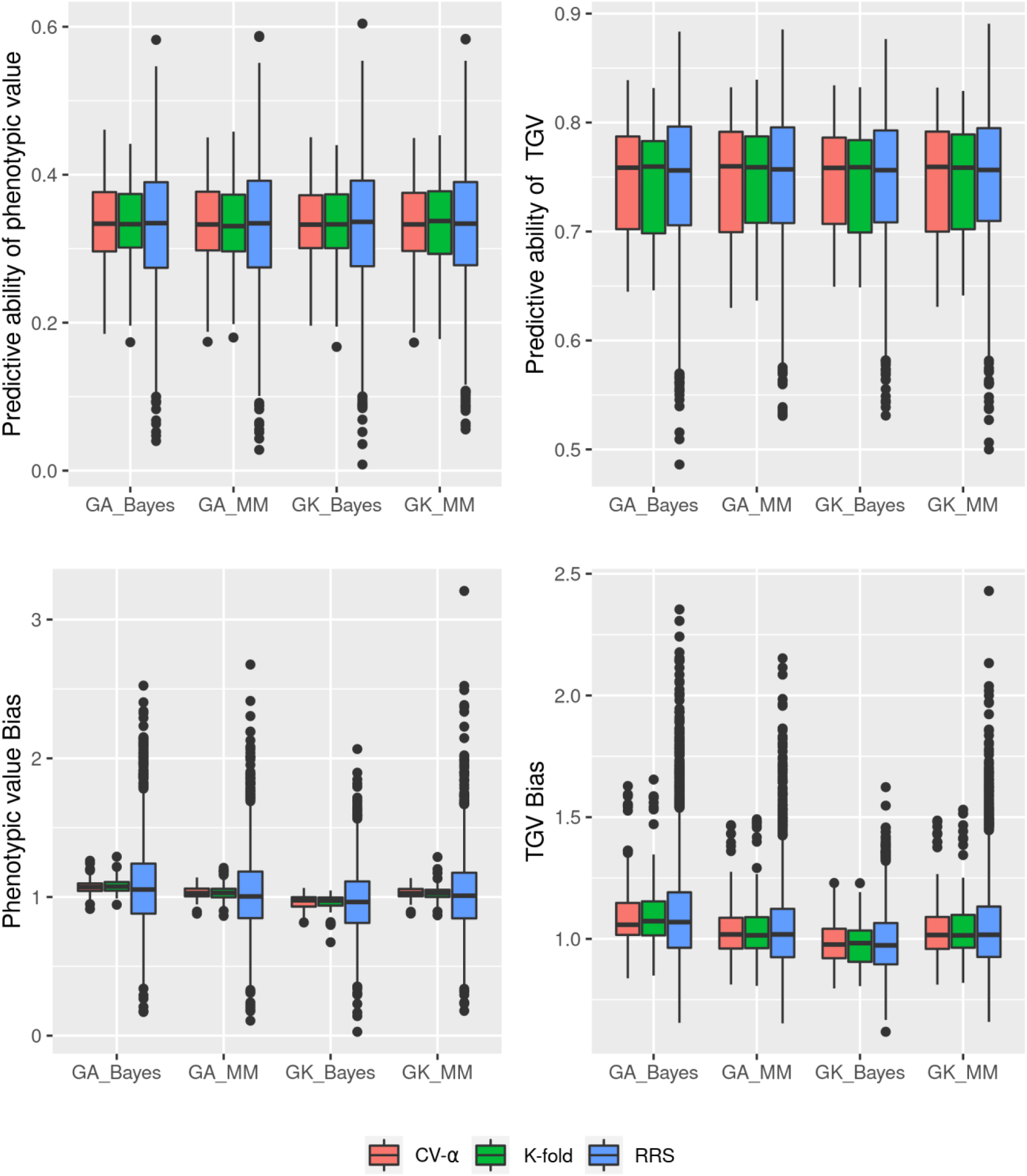
Predictive ability (PA) and bias for TGV and phenotypic value for three validation schemes (CV-α, K-fold, and RRS) and four genomic prediction models (scenarios).

**Figure 4.**
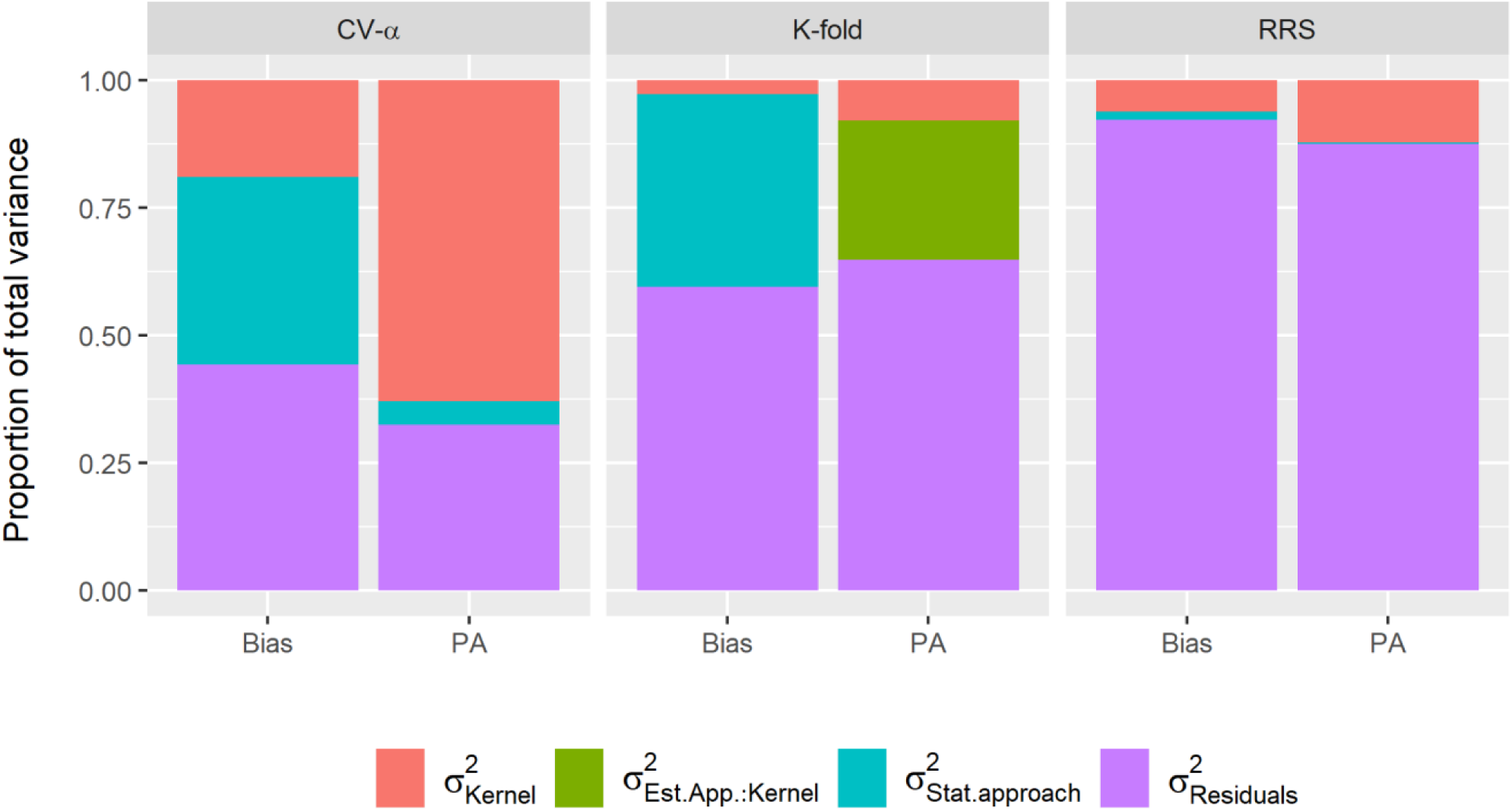
The proportion of total variance decomposed into effects of the kernel, statistical approach, the interaction between the kernel and statistical approach, and residual for bias and predictive ability (PA) applied in three cross-validation schemes (CV-α, K-fold, RRS).

For PA and bias, TGV, and phenotypic value, in terms of mean and standard deviation, the three validation methods do not differ among them (Table 1), except for phenotypic bias for RRS. On the other hand, when we considered mean squared error (MSE) and coefficient of variation (CV), CV-α showed the lowest CV for all scenarios evaluated, when compared with RRS and K-fold.

**Table 1.**
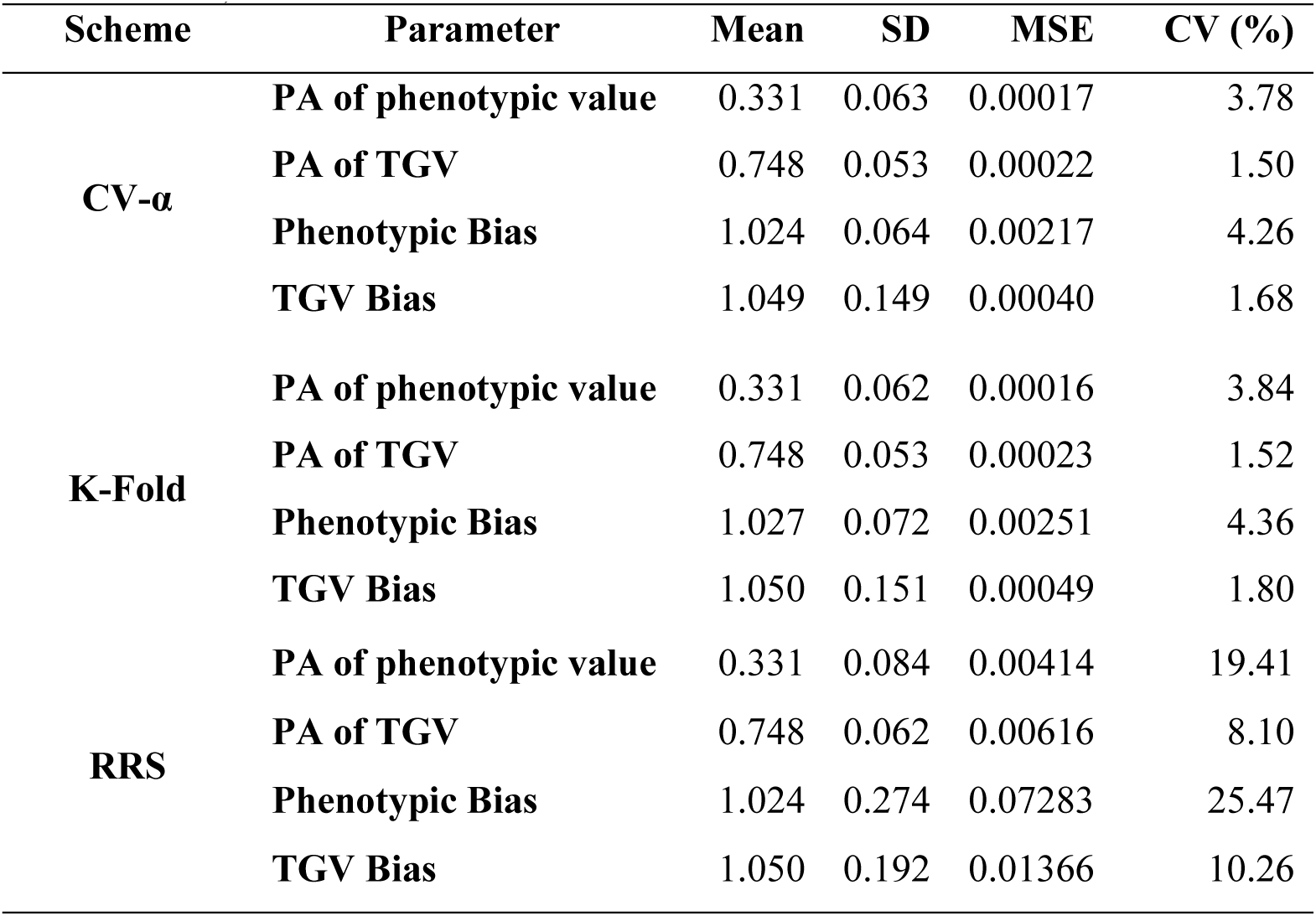
Averaged of 25 simulated datasets for mean, standard deviation (SD), mean squared error (MSE), and coefficient of variation (CV) for predictive ability (PA) and bias for three CV schemes (CV-α, K-fold, and RRS)

### Proof of concept

For the maize dataset, PA and bias showed similar mean values for all validation methods. In terms of SD, K-fold, and CV-α presented similar performance and were lower than RRS (Table 2).

**Table 2.**
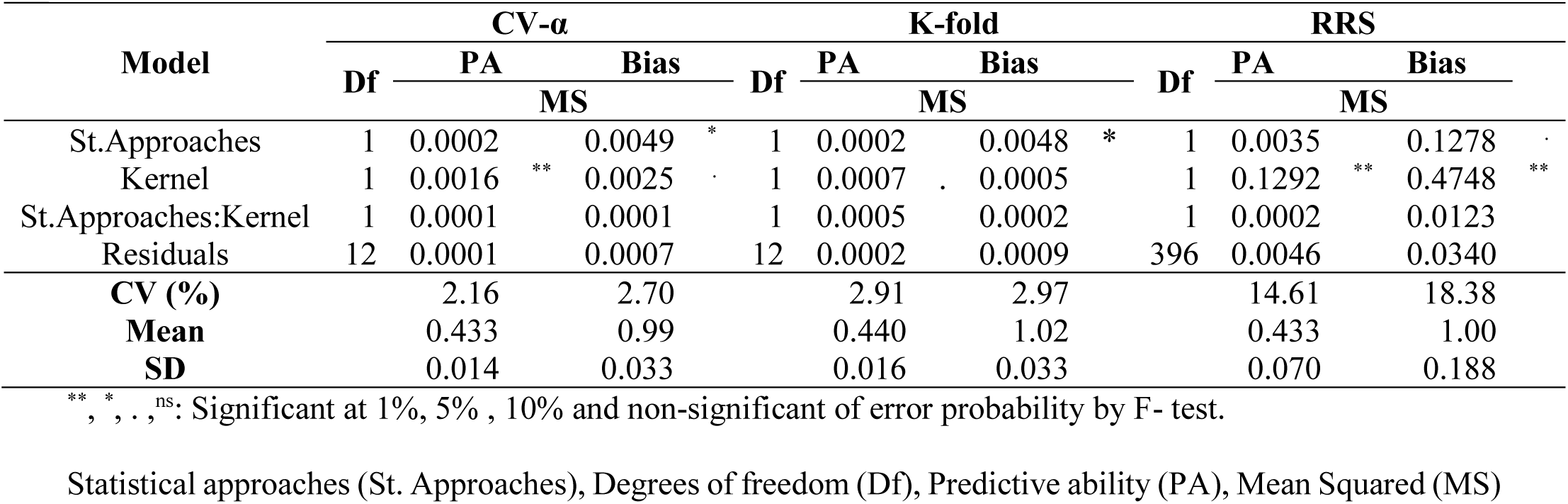
Summary of ANOVA, mean, standard deviation (SD), and coefficient of variation (CV) for three validation schemes (CV-α, K-fold, and RRS) for predictive ability (PA) and bias

For mean squared error and coefficient of variation, CV-α presented lower values than K-fold and RRS. The coefficient of variation for K-fold was 34.70% and 10% higher than CV-α for PA and bias, respectively (Table 2).

We applied the CV-α to validate two statistical approaches (Bayesian and Mixed models) and two types of kernels (Additive and Gaussian kernel) for genomic prediction models (Table 3). For this, we applied a two-way ANOVA, and it was observed significative effects for types of the kernel for predictive ability and bias. Gaussian kernel (**K**) presented higher PA (0.44) than **G**_**a**_ (0.42) and lower bias (1.00 and 0.98, for **K** and **G**_**a**_, respectively). For the type of two statistical approaches, the Bayesian reveals a more biased estimation (0.98) when compared with GBLUP (1.01) (Table 3).

**Table 3.**
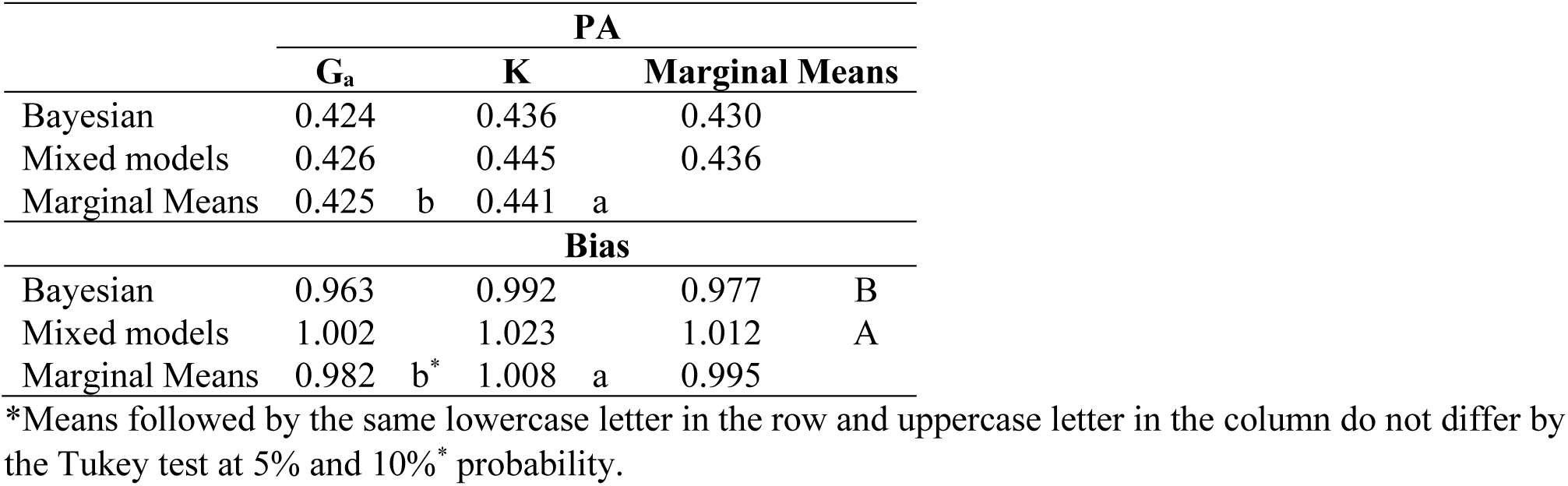
Means, marginal means, and Tukey’s test for the type of kernels and statistical approaches for predictive ability (PA) and bias

|  | PA |  |  |  |  |
| --- | --- | --- | --- | --- | --- |
|  | G<sub>a</sub> |  | K |  | Marginal Means |
| Bayesian | 0.424 |  | 0.436 |  | 0.430 |
| Mixed models | 0.426 |  | 0.445 |  | 0.436 |
| Marginal Means | 0.425 | b | 0.441 | a |  |
|  | Bias |  |  |  |  |
| Bayesian | 0.963 |  | 0.992 |  | 0.977 B |
| Mixed models | 1.002 |  | 1.023 |  | 1.012 A |
| Marginal Means | 0.982 | b* | 1.008 | a | 0.995 |

\*Means followed by the same lowercase letter in the row and uppercase letter in the column do not differ by the Tukey test at 5% and 10% \* probability.

We can note that the proportion of phenotypic variance explained variation by each source of variation vary across validation schemes (Figure 3). PA and bias had similar performance across CV schemes for residual variance but vary for other variances. The RRS presented higher residual variance and lower variances due to model effects. For the interaction, K-fold showed higher values for PA. CV-α presented lower proportions of residual variances and higher variance due to the kernel and statistical approaches effects.

## Discussion

The main advantages of considering the α-design instead of the balanced incomplete block design (BICV) are the flexibility regarding the number of treatments and folds (Singh e Bhatia 2017), reduce the concurrence between pairs of treatments, increase the quality of estimates (Fuchs e Krautenbacher 2016) and residuals independence, allowing further multiple comparison analyses. The α-design is widely used in plant breeding experiments as well as its ANOVA (Alam et al. 2017; Ta et al. 2018; Galic et al. 2019). Based on this, in the context of genomic prediction, the flexibility of the CV-α is a good alternative to compare genomic selection models.

Our results reveal that CV-α reduces the concurrence between pairs of treatments (genotypes) in the same fold across replicates and its standard deviation when compared with the K-fold scheme (Figure 2). The concurrence of any two treatments causes dependence among folds, and comparative tests become less precise. Thus, the CV-α designs fold and replicates with few or non-concurrence across folds, generating a more independent and better scheme for composing training and validation sets in a genomic prediction context.

Comparison between CV-α, K-fold, and RRS must be pondered since they have a different level of averaging and different numbers of replicates compared with RRS, although CV-α and K-fold are equivalent (Wong 2015). RRS showed a higher number of outliers, probability as results of the different levels of average. However, it is an internal procedure for the method. The strategy to divide folds and replicates according to the alpha-lattice design, as we suggest into CV-α, permits we consider as replicate level mean, similar to replicate the effect in the alpha-lattice design.

Moreover, the RRS showed a large variation in the estimates for PA and, especially, for prediction bias. We expected values for bias around 1.0. However, the RRS showed several values overtaking 0.5 and 1.5, which shows a considerable inflation/deflation on the estimates. These results indicate that RRS is a less accurate method, mainly when we use few replicates.

Estimates more accurate combined with few replicates to run a CV scheme is desirable, especially when we consider a large number of genotypes, which is common in plant and animal breeding. In these cases, to compute the inverse matrix, the genomic relationship matrix is a challenge, and several studies have been aiming this (Misztal et al. 2014; Misztal 2016). Based on this, CV-α is a good alternative to design CV schemes and has a more precise estimative in a case where the number of replicates is a limitation.

The simulated and real datasets reveal that CV-α had a similar performance to K-fold and RRS when compared in terms of mean and standard deviation for predictive ability and bias. On the other hand, when we consider in terms of MSE and coefficient of variation, CV-α has better performance due to higher independence across replicates.

Traditionally, in genomic prediction studies, model comparison and selection are based on subjective methods such as mean and standard deviation without a comparative test. Some studies also considered ANOVA and other statistical tests. Although due to assumption unfulfilled regarding residuals independence and our results, this is not be recommended. CV-α reveals the lesser occurrence of pairs of genotypes in the same fold across replicates, causing a more precise estimative. The CV-α methodology consists of applying α(0,1) lattice design to design the folds across replicates, and because of this, it allows post hoc test to model comparison.

The results above indicate that CV-α had a more precise estimative trough the reduction of coefficient of variation, and the variance components were better discriminated across the factors in the two-way ANOVA. It reveals how the impact of folds design across each replicate shift the proportion of the total variation explained by each model factor reducing the residual variance. Furthermore, the ANOVA test using RRS and K-fold to compare the performance of different models can produce mistake conclusions, since the estimative of variance components load bias. Therefore, CV-α allows determining how much variation each model factor has and compares different genomic selection models based on the ANOVA test and posthoc test. Furthermore, the use of CV-α does not imply any additional computer cost or complexity in the validation process of model selection.

As proof of concepts, we applied the proposed methodology to exemplify model selection. For the simulated and maize dataset, both do not show considerable differences across approaches (GBLUP and Bayesian) and kernel type (Additive genomic and Gaussian kernel) for predictive ability. Although, for the maize dataset, the use of **K** kernel showed higher predictive ability than **G**_**a**_. This result is expected since the **K** kernel captures additive and non-additive effects (Heslot et al. 2012). For bias, mixed models showed less biased results. Although, comparison among these models is not the focus of these studies since they have already been extensively studied (Chen et al. 2014; Gota e Gianola 2014; Cuevas et al. 2017).

In the context of genomic prediction studies, there are other ways to design training and validation sets. The CV-α may be expanded for these cases to better designing training and test sets across replicates and environments, such as CV1 and CV2 schemes (Burgueño et al. 2012) and other multi-environment and multi-trait studies. Also, the CV-α may be applied in any other cross-validation studies to select models and verify as the model factors behave according to the different sources of variation.

## Conclusion

This study showed that the CV-α method is a good alternative to design cross-validations folds and replicates, mainly when researchers want to compare genomic prediction models, increasing precision in the model estimative, and to unravel the model factors impact in the total variation. Even though there were no differences in the mean and standard deviation for predictive ability and bias, our proposal was more accurate in terms of the mean squared error and coefficient of variation. Another advantage of CV-α is that it does not require any additional cost regarding computing demand or complexity. Furthermore, CV-α allows using the non-subjective methods to compare models and factors, through ANOVA and other multiple comparison tests, such as Tukey and Scott-Knott.

## Abbreviations

CV: cross-validation
CV-α: cross-validation alpha-based design
GP: Genomic prediction
RRS: Repeated Random Subsampling
TGV: true genetic value

## Acknowledgment

This study was financed in part by the Coordenação de Aperfeiçoamento de Pessoal de Nível Superior - Brasil (CAPES) - Finance Code 001, Conselho Nacional de Desenvolvimento Científico e Tecnológico (CNPq).

